# Seminal plasma inhibits Chlamydia trachomatis infection *in vitro*, and may have consequences on mucosal immunity

**DOI:** 10.1101/2023.10.10.561648

**Authors:** Louis Reot, Cindy Adapen, Claude Cannou, Natalia Nunez, Sabrine Lakoum, Camille Pimienta, Laetitia Lacroix, Olivier Binois, Nelly Frydman, Marie-Thérèse Nugeyre, Roger Le Grand, Elisabeth Menu

**Affiliations:** Université Paris-Saclay, Inserm, Commissariat à l’Energie Atomique et aux Energies Alternatives (CEA), Center for Immunology of Viral, Auto-Immune, Hematological and Bacterial Diseases [IMVA-HB/Infectious Disease Models and Innovative Therapies (IDMIT)], Fontenay-aux-Roses, France; Institut Pasteur, Université Paris Cité, Mucosal Immunity and Sexually Transmitted Infection Control (MISTIC) Group, Department of Virology, Paris, France; Life&Soft, Fontenay-aux-Roses, France; Université Paris-Saclay, Assistance Publique Hôpitaux de Paris, Hôpital Antoine Béclère, Service de Biologie de la Reproduction CECOS, Clamart, France

**Keywords:** STI, *C. trachomatis*, seminal plasma, inflammation, female mucosa, neutrophil

## Abstract

Seminal plasma (SP) is the main vector of *C. trachomatis* (CT) during heterosexual transmission from male to female. It has immunomodulatory properties and impacts the susceptibility to HIV-1 infection, but its role has not been explored during CT infection. In the female reproductive tract (FRT), CT infection induces cytokine production and neutrophil recruitment. The role of neutrophils during CT infection is partially described, they could be at the origin of the pathology observed during CT infection. During this study, we developed an experimental *in vitro* model to characterize the impact of CT infection and SP on endocervical epithelial cell immune response in the FRT. We also studied the impact of the epithelial cell response on neutrophil phenotype and functions. We showed that the production by epithelial cells of pro-inflammatory cytokines increased during CT infection. Moreover, the pool of SP as well as individuals SP inhibited CT infection in a dose-dependent manner. The pool of SP inhibited cytokine production in a dose-dependent manner. The pool of SP altered gene expression profiles of infected cells. The culture supernatants of cells infected or not with CT, in presence or not of the pool of SP, had an impact on neutrophil phenotype and functions: they affected markers of neutrophil maturation, activation and adhesion capacity, as well as the survival, ROS production and phagocytosis ability. This study proposes a novel approach to study the impact of the environment on the phenotype and functions of neutrophils in the FRT. It highlights the impact of the factors of the FRT environment, in particular SP and CT infection, on the mucosal inflammation and the need to take into account the SP component while studying sexually transmitted infections during heterosexual transmission from male to female.

## Introduction

Sexually transmitted infections (STI) represent a major public health issue, with a growing incidence of several STI, despite the existence of different treatments. Over the last years, *Chlamydia trachomatis* (CT) infection cases increased worldwide, especially in young people^1,2^. CT is a Gram-negative bacterium that infects preferentially epithelial cells. Serovars D to K are responsible for urogenital infections. Most of the cases of CT infection are asymptomatic, and the infected population represent a reservoir for STI spreading, so there is a real need for prevention methods. One of the key issues to develop new preventive strategies is to understand more deeply the role of the local environment during CT acquisition.

In the female reproductive tract (FRT), CT infection results in a pro-inflammatory cytokine and chemokine response that leads to the recruitment of innate, and later adaptive immune cells, allowing the control of the infection^3–5^. Persistence of the pathogen results in chronic inflammation and can lead to collateral genital tract tissue damage^6,7^. Polymorphonuclear leucocytes (PMN) from the blood are the first immune cells recruited to the site of CT infection. Through various antimicrobial activities (degranulation, phagocytosis, reactive oxygen species (ROS) and neutrophil extracellular traps (NET) production …), they allow pathogen clearance. However, the role of neutrophils during CT infection is not fully understood. They could induce the immune pathology during CT infection^8^: in an *in vitro* study, CT enhanced neutrophil responses, characterized by a recruitment of neutrophils to the site of infection, their activation and prolonged survival^9^. However, in other experimental settings, neutrophil responses are counteracted, and neutrophils are paralyzed by CT^10^. Moreover, in *in vivo* mouse models, neutrophils can contribute to antibody-mediated protection to CT infection^11^, or on the contrary, drive CT associated pathology, without reducing the bacterial burden^8^. These studies highlight the need to better characterize the role of those cells during CT infection.

During heterosexual transmission from male to female, the main vector of STI pathogens in the FRT is the seminal plasma (SP)^12,13^. Very little is known about the role of seminal plasma during CT induced pathology. SP, the acellular fraction of the semen, is composed of various proteins, including pro and anti-inflammatory cytokines^14^. It is known to induce an inflammatory reaction in the FRT, leading to cytokine production and the recruitment of neutrophils from the blood^15,16^. The inflammation induced by SP exposure in the FRT has been linked to the regulation of infectious diseases, notably modifying the risk for HIV-1 acquisition^17,18^. SP also has strong antimicrobial properties conferred by proteolytic cleavage of semenogelins (the main protein of human semen coagulum), displaying bactericidal activity against various Gram-positive and Gram-negative bacteria^19^. However, even if SP has been shown to modulate STI susceptibility and inflammation at the level of the FRT^20^, the impact of SP is poorly investigated during CT infection.

In this study, we aimed at characterizing the impact of CT infection in presence or in absence of SP on epithelial cell response at the level of the FRT mucosa. We also investigated the impact of the epithelial cell response on the phenotype and functions of neutrophils. Using an *in vitro* model, we found that SP inhibited CT infection and modified the inflammation induced by the infection, in particular the cytokine profile of CT infected epithelial cells. We also show that the local environment could have an impact on local neutrophil phenotype and functions.

## Material and Methods

### Seminal plasma collection

Human seminal plasmas were obtained from the “*Centre d’étude et de conservation des œufs et du sperme humains*” (CECOS) at Antoine Béclère Hospital. Semen was collected from patients that were involved in *in vitro* fertilization protocol. All methods to recruit participants and collect biological samples were carried out in accordance with relevant guidelines and regulations. Participants gave their consent for the use for medical research purposes of the samples. They were informed of the study and gave no objection for the use of their samples and data for the study.

The study was defined as a non-interventional study by French Ethical Committee ‘Comité de Protection des personnes’ (n° 2014/42NICB). The study complies with the methodology reference MR-004 set out by the French data protection authority (‘Commission Nationale de l’Informatique et des Libertés’ CNIL) for which Institut Pasteur Paris made a statement compliance with the CNIL (n° 2214728v0). The study was registered on the public directory of the health data hub HDH (N° F20220617111005).

All semens had a negative serology for HBV, HCV, HIV and Syphilis, and were negatively tested for Chlamydia and other bacterial infections by spermaculture. SP were obtained after semen centrifugation on gradient density.

SP from about 100 patients were pooled, aliquoted and characterized for cytokine concentration. Thirty-two individual SP were also stored.

### Cell line culture

A2EN is a human endocervical epithelial cell line, generated in the laboratory of Dr A. Quayle from primary epithelial cells isolated from endocervical explant, and immortalized with human papilloma virus E6 and E7^21^. A2EN cells (Kerafast) were grown in phenol red-free serum-free medium (EpiLife; Cascade Biologics) with an Epilife Defined Growth Supplement (EDGS; Gibco), 1% Penicillin/Streptomycin (PS) and 0.004M of CaCl_2_ (Sigma Aldrich). The cells were grown at 37°C with 5% CO_2_. High molecular weight (HMW) poly(I:C) (100 µg/mL) was used as a positive control to stimulate A2EN cell cytokine production.

### *C. trachomatis* infection

*Chlamydia trachomatis* serovar D (D/UW3/Cx) was obtained from Statens Serum Institut (Copenhagen, Dr Follmann laboratory). A2EN cells were seeded in EpiLife medium with EDGS, CaCl_2_ and PS in 24 well plate at 5.10^4^ cells/well. After 96h, confluent A2EN cells were washed, and fresh Epilife medium without PS containing, or not, various dilutions of SP (1/10, 1/50, 1/100 or 1/500) was added. The cells were immediately infected with CT svD at different multiplicity of infection (MOI) (12, 25 or 50) by centrifugation (700 g) for one hour. Then, culture medium was removed and DMEM with 10% FCS was added. The cells were incubated for 24h at 37°C with 5% CO2. After 24h, pH measurement (Fisherbrand™ pH Indicator, 6.4-8) and viability test (CellTiter®) were performed. Supernatants were collected and frozen at −80°C after 0.2μm filtration. For the determination of the percentage of infection, cells were fixed and stained using a rabbit anti-chlamydia heat shock protein 60 (Hsp60) antibody (provided by A. Subtil, Institut Pasteur, France) diluted at 1:2000 in PBS + 1% BSA for 1h. Then, cells were treated with anti IgG Rabbit – AF488 (1/500), DAPI and Phalloidin for 1h. Inclusions were quantified in infected cells using an inverted microscope (Zeiss® Axiovert 25). The percentage of infection was determined by dividing the number of inclusions containing cells by the total number of cells.

### RNA extraction and sequencing

At different timepoints, cells were lysed and lysates stored at −80°C for RNA extraction. RNA was extracted using NucleoSpin RNA XS kit (Macherey Nagel) according to manufacturer instructions. RNA was quantified using QuBit RNA HS kit (ThermoFisher), and a quality check was performed on the Agilent TapeStation system. A total of 1000 ng of RNA per sample was denatured at 65°C and retrotranscribed by a strand-switching technique using Maxima H Minus Reverse Transcriptase (ThermoFisher, USA) to synthesize a double stranded cDNA. PCR, barcode, and adapter attachment were performed according to cDNA-PCR Sequencing Kit (SQL-PCB109, Oxford Nanopore Technologies, Oxford, UK). Samples were quantified using QuBit dsDNA HS (ThermoFisher, USA) kit before loading on R9.4.1 Flow cells using the GridION instrument (Minknow version 21.11.7).

### Transcriptome Analysis

Sequence reads were converted into FASTQ files. Reads under 300 bp or with a quality score under 9 were discarded. The remaining reads were aligned on the human GRCh38.p13 and *C. trachomatis* D/UW-3/CX strain transcriptome of reference (GeneBank assembly accession numbers GCA_000001405 and GCA_000008725 respectively) using minimap2^22,23^ version 2.24. To quantify transcripts, the resulting alignments were given to Salmon version 1.8.0^24^. To explore differentially expressed genes, replicates count data were used on DESeq2 version 1.32.0^25^ in one single DEseq model. Gene set enrichment analysis with both upregulated and downregulated genes Log2FC>1.5 or Log2FC<1.5, respectively was performed using Enrichr, a web server enrichment analysis tool^26,27^, and BioPlanet 2019 database for cellular and signaling pathway analysis^28^.

### Cytokine and Chemokine quantification

Pro- and anti-inflammatory cytokines were measured in the A2EN cell culture supernatants and in the seminal plasma by a 25plex assay for the detection of: IL-1β, IL-1RA, IL-2, IL-2R, IL-4, IL-5, IL-6, IL-7, IL-8, IL-10, IL-12 (p40/p70), IL-13, IL-15, IL-17, TNFα, IFNα, IFNγ, GM-CSF, CCL3, CCL4, CXCL10, CXCL9, CCL11, CCL5 and CCL2 (Human cytokine magnetic 25-plex panel; Life technologies).

TGF-β1, 2 and 3 were detected with another multiplex assay (MILLIPLEX MAP TGFß Magnetic Bead 3 Plex Kit; Merck Millipore).

### Neutrophil isolation, phenotype and functional analysis

Human neutrophils were isolated from human blood, obtained from blood donors at *Etablissement Francais du Sang* (EFS, France; C CPSL UNT-N°13/EFS/101), using the EasySep™ Direct Human Neutrophil Isolation Kit according to the manufacturer’s instructions (StemCell Tech, Vancouver, BC, Canada). Each sorting resulted in more than 90% neutrophils, confirmed by phenotype analysis. 2.10^5^ neutrophils were incubated at 37°C or at 4°C (only for the phagocytosis assay) in a 96-well plate containing an equal volume of RPMI with 10% FCS and of A2EN cell supernatants or DMEM with 10% FCS. Phenotype and functional analysis were performed at various time post-incubation.

Survival of the neutrophils was followed at 2h, 6h, 24h and 48h using an apoptosis detection kit with PE-Annexin V and 7-AAD (Biolegend), according to the manufacturer instructions. At 2h post-incubation, neutrophils were tested for ROS production and phagocytosis capacity.

For the phagocytosis assay, pHrodo™ Red *E. coli* BioParticles™ Conjugate for phagocytosis (Invitrogen™) were added to the neutrophils at a 1:1 ratio for 5min, then cells were washed and fixed using 1% PFA until flow cytometry analysis.

For the ROS production assay, neutrophils were washed and incubated at 37°C for 90min in PBS containing 10µM Luminol (Sigma-Aldrich). PMA (0,1ng/µL, Sigma-Aldrich) was added 0min or 50min after the addition of the Luminol. Luminescence was followed every min using a multimode microplate reader, the Spark 10M (TECAN; Switzerland). The phenotype of the cells was analysed after 6h of incubation, after staining of the neutrophils for 10min at 4°C with the antibodies listed in Table 1. A ZE5 flux cytometer was used (Biorad) with Everest (Biorad) and FlowJo (Tristar, USA) software packages for the flow cytometry analysis. A representative image of the gating strategy is illustrated in Supplementary Figure 1.

**Table 1:**
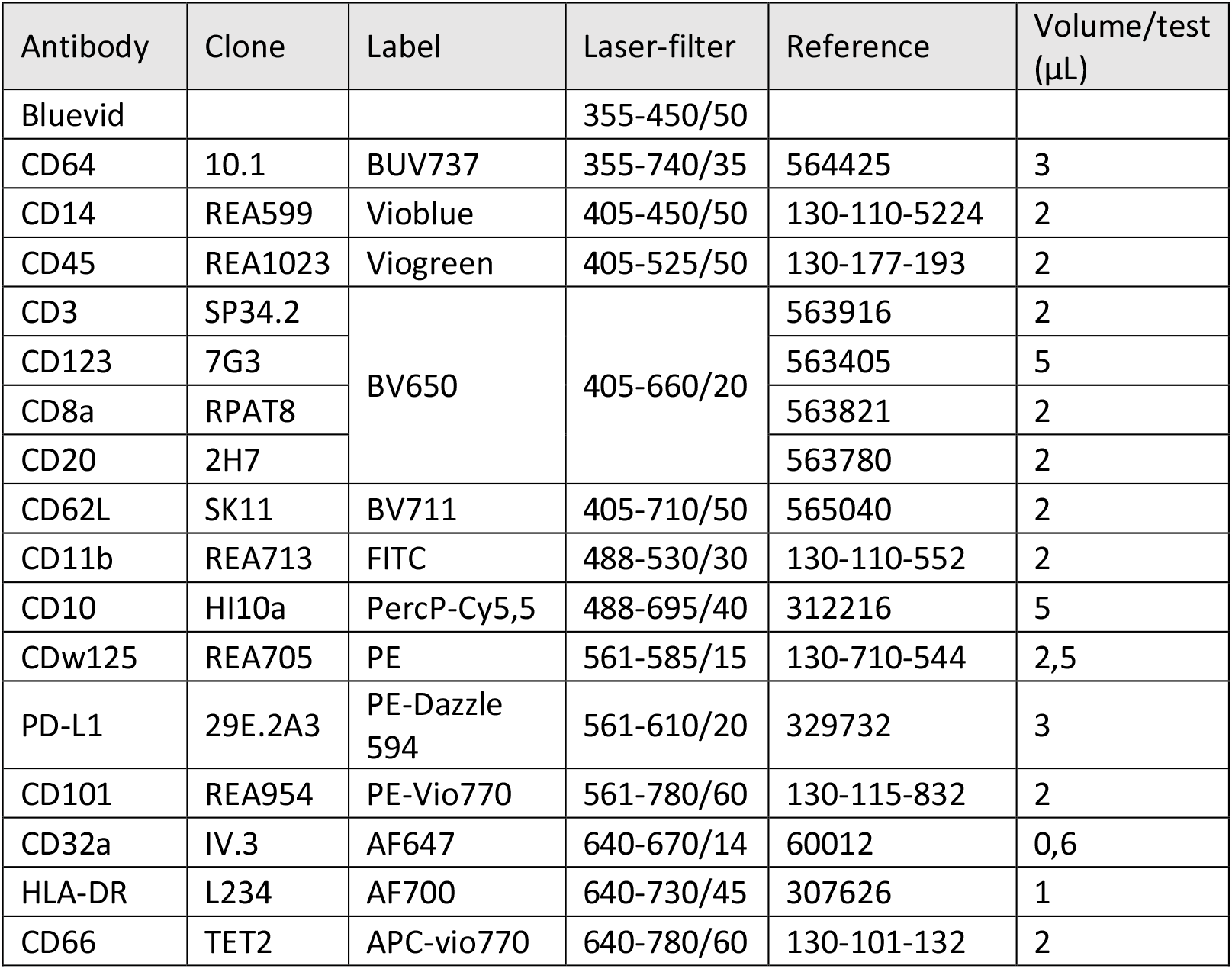
Antibody panel used to characterize neutrophil populations.

| Antibody | Clone | Label | Laser-filter | Reference | Volume/test (µL) |
| --- | --- | --- | --- | --- | --- |
| Bluevid |  |  | 355-450/50 |  |  |
| CD64 | 10.1 | BUV737 | 355-740/35 | 564425 | 3 |
| CD14 | REA599 | Vioblue | 405-450/50 | 130-110-5224 | 2 |
| CD45 | REA1023 | Viogreen | 405-525/50 | 130-177-193 | 2 |
| CD3 | SP34.2 | BV650 | 405-660/20 | 563916 | 2 |
| CD123 | 7G3 |  |  | 563405 | 5 |
| CD8a | RPAT8 |  |  | 563821 | 2 |
| CD20 | 2H7 |  |  | 563780 | 2 |
| CD62L | SK11 | BV711 | 405-710/50 | 565040 | 2 |
| CD11b | REA713 | FITC | 488-530/30 | 130-110-552 | 2 |
| CD10 | HI10a | PercP-Cy5,5 | 488-695/40 | 312216 | 5 |
| CDw125 | REA705 | PE | 561-585/15 | 130-710-544 | 2,5 |
| PD-L1 | 29E.2A3 | PE-Dazzle 594 | 561-610/20 | 329732 | 3 |
| CD101 | REA954 | PE-Vio770 | 561-780/60 | 130-115-832 | 2 |
| CD32a | IV.3 | AF647 | 640-670/14 | 60012 | 0,6 |
| HLA-DR | L234 | AF700 | 640-730/45 | 307626 | 1 |
| CD66 | TET2 | APC-vio770 | 640-780/60 | 130-101-132 | 2 |

### Statistical analysis

GraphPad prism software version 9 for windows (GraphPad Software, La Jolla California USA, www.graphpad.com) was used for graphical representations and area under the curve (AUC) calculation. Two-way ANOVA test with p values adjustment with Tukey’s test was used for statistical analysis.

## Results

### Seminal plasma composition

The pool of SP was characterized in terms of cytokine/chemokine concentrations (Figure 1). The pool of SP had a moderate inflammatory profile, with some pro-inflammatory cytokines expressed (IL-8, CCL2, CXCL9) but also anti-inflammatory cytokines (TGF-β, IL-1RA). As expected^29^, TGF-β1, mβ3 and –β2, respectively, were the most prevalent cytokines in seminal plasma.

**Figure 1:**
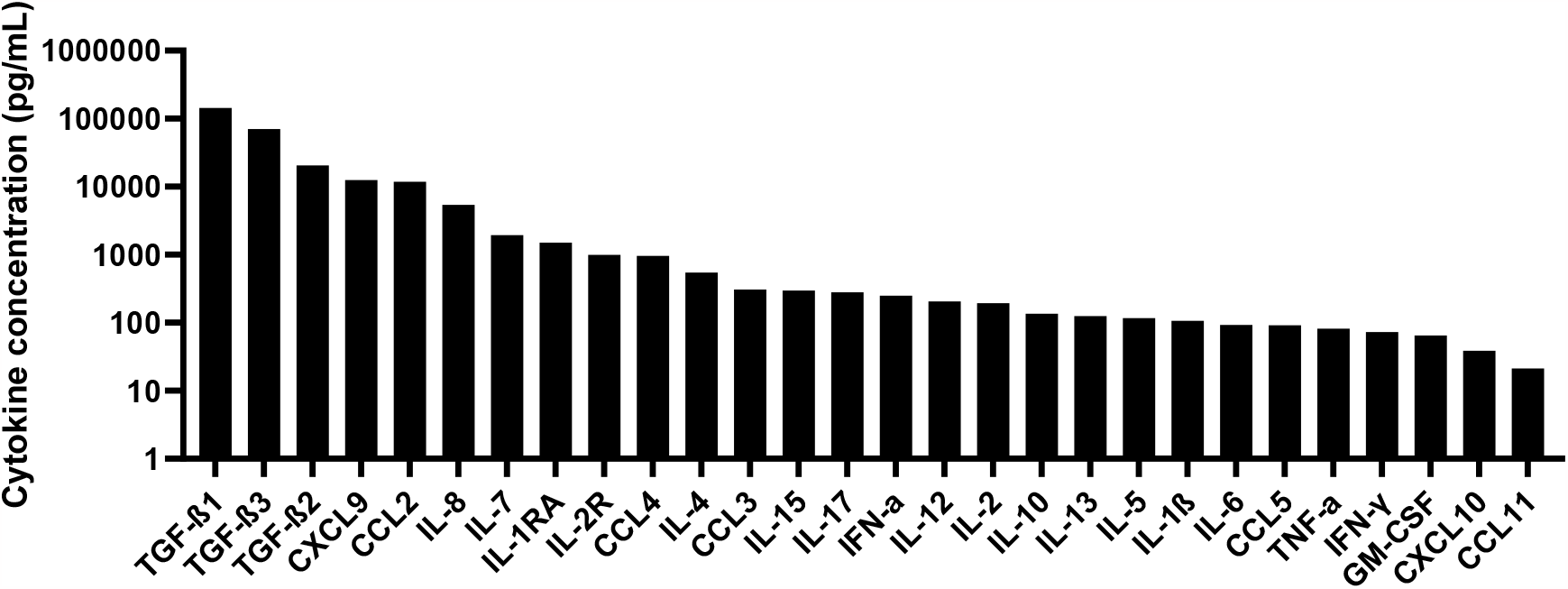
Cytokine characterization of the pool of SP. Seminal plasma from about 100 patients were pooled and the cytokine concentrations of the pool were quantified by Luminex®.

### Seminal plasma inhibited *Chlamydia trachomatis* infection of A2EN cells

The effect of the pool of SP was examined on A2EN cells infected with CT. Since pure SP could affect the viability of epithelial cells *in vitro*, we tested several dilutions of SP in PBS, starting with a 10-fold dilution, to approximate the physiological situation in the FRT after intercourse^30^. No effect was detected on cell viability: the pH of A2EN supernatants and the metabolic activity of the cells were similar in all the conditions tested (Figure 2A, 2B). We then determined the percentage of CT infection in the various conditions by immunofluorescence. A representative image is illustrated in Figure 2C. We quantified the cells that had at least one inclusion and determined that the presence of SP drastically reduced the percentage of CT infected cells analyzed 24h post-infection (Figure 2D). This effect was dose dependent: the percentage of infection decreased with the SP concentration. The 10-fold dilution of SP was the most potent to reduce the percentage of CT infection, with 10-fold less infected cells compared to the condition without SP. No significant difference in the size of the inclusions was observed between the various experimental conditions (Supplementary Figure 2). We then tested if this effect could be observed using different experimental conditions. The results represented in Supplementary Figure 3 show that SP also inhibited CT infection at higher MOI or at 48h post-infection. Uninfected A2EN cells produced various cytokines including CXCL10, IL-6, GM-CSF and a high concentration of IL-8 (Figure 3A). Stimulation with poly(I:C), strongly increased A2EN cell production of several proinflammatory cytokines, including CCL5, CCL3, CXCL10 and IL-6 (Fig 3B). CT infection induced an increase in inflammatory cytokines concentration (IL-6, GM-CSF and CXCL10), but lower than poly(I:C). In contrast, when cells were exposed to SP alone, a decrease in the concentrations of those same cytokines was observed in a dose dependent manner (Figure 3B). When A2EN cells were exposed to CT in presence of SP, the cytokine profile was dependent on the dilution of SP: the effect of SP predominated for low dilutions, with a decrease in cytokine concentration. On the contrary, for high dilutions, the effect of CT predominated, with an increase in inflammatory cytokine concentration. In conclusion, SP inhibited CT infection and decreased the inflammation induced by CT.

**Figure 2:**
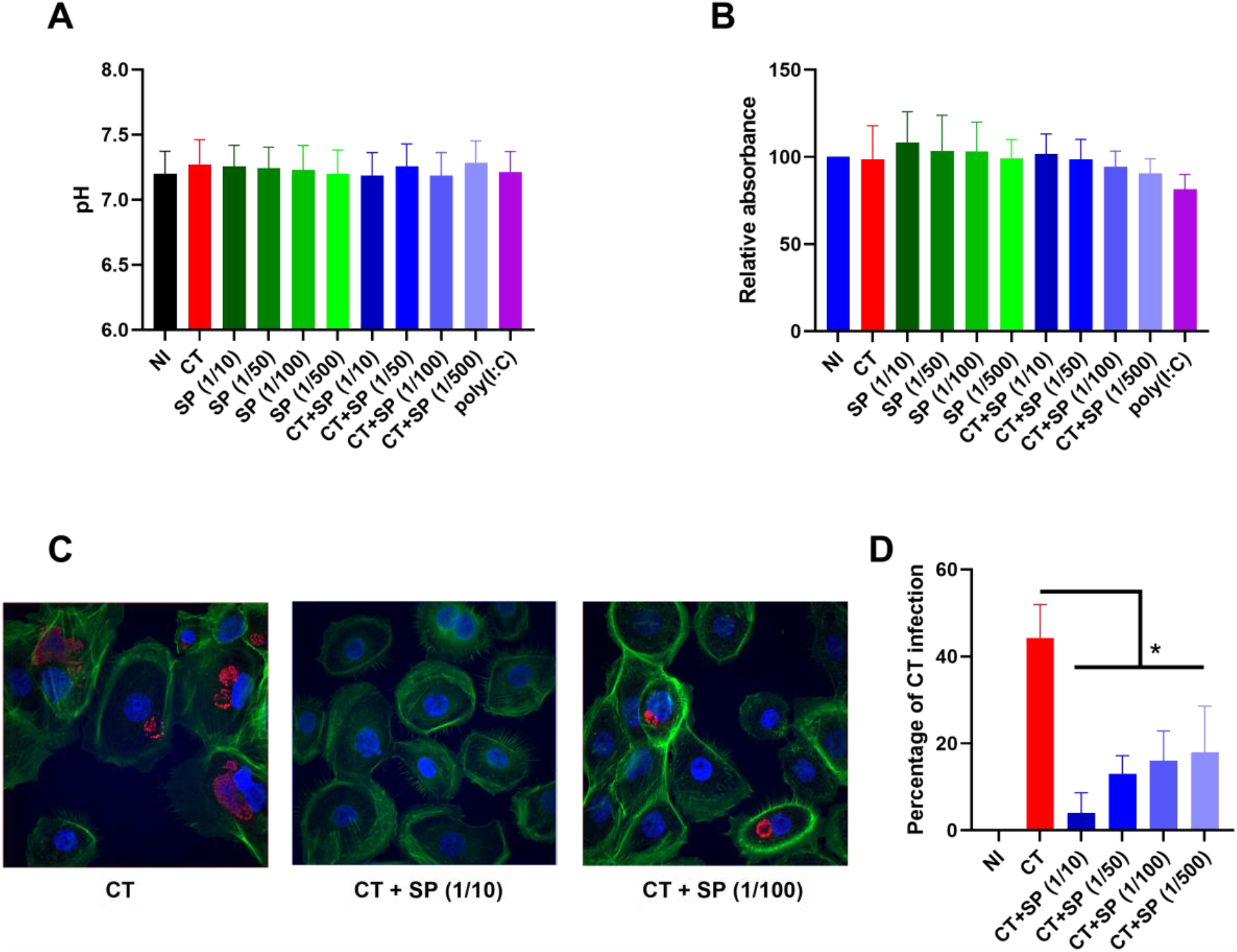
Impact of the pool of SP on CT infection of A2EN cells. Cells were infected or not with CT at a MOI of 12, in the presence or not of different dilutions of the pool of SP, for 24h (n=4). Poly(I:C) was used as a positive control. (A) pH was measured in the supernatants. (B) A2EN viability was evaluated using CellTiter and the relative absorbance was measured. (C) The percentage of infection was determined by immunofluorescence. Actin filaments in the cytoplasm are in green (Phalloidin), the nucleus in blue (DAPI), and the CT inclusions in red (Hsp60 antibody). (D) Percentage of CT infection (?number of infected cells/total number of cells] x 100) at 24hpi by quantification of the inclusion in A2EN infected cells. The asterisk indicates a significant difference by T-test (*p ≤0.05).

**Figure 3:**
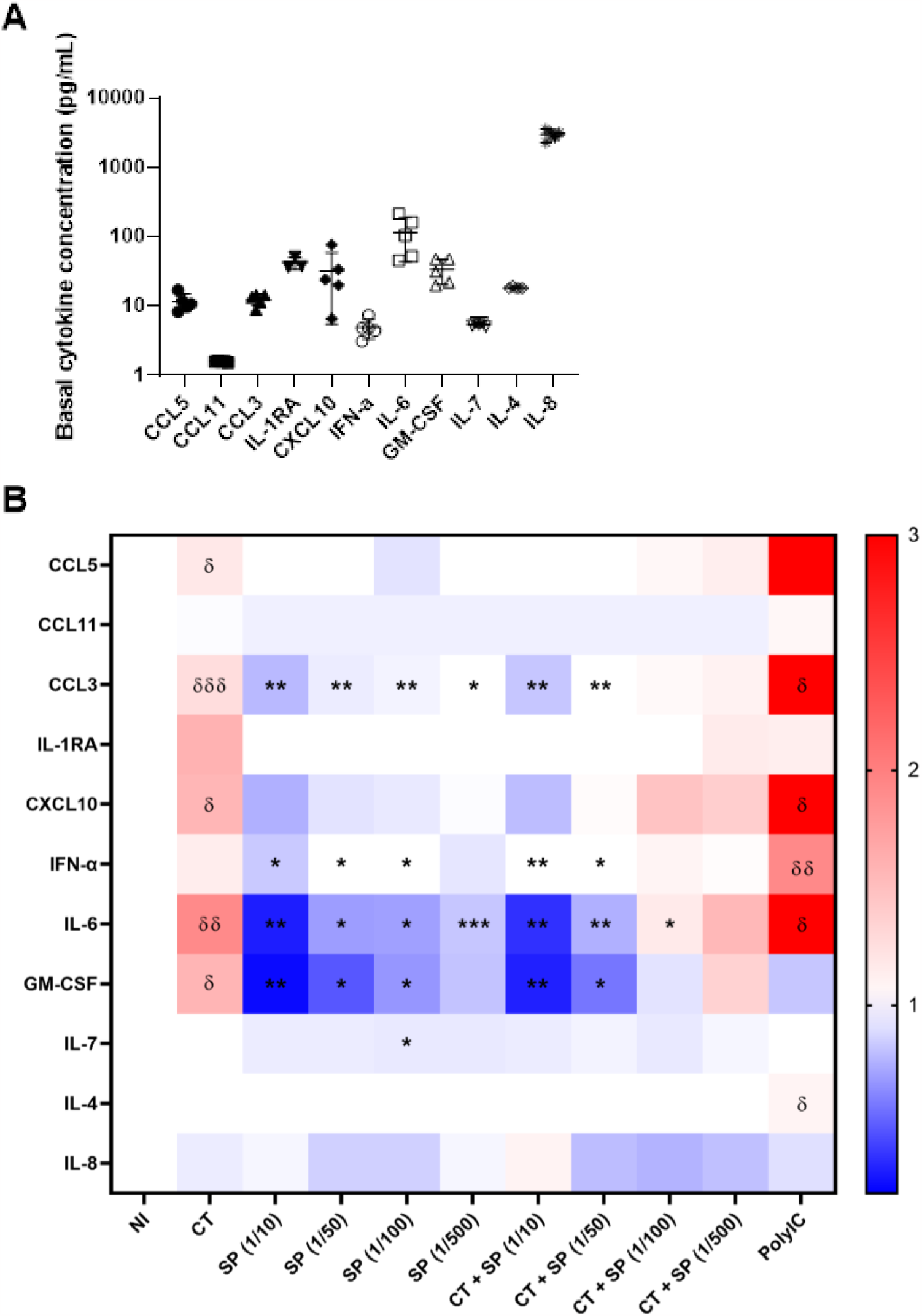
Impact of the pool of SP on cytokine expression by A2EN cells infected with CT. Cells were infected or not with CT at a MOI of 12 for 24h, in the presence or not of different dilutions of the pool of SP (n=5). Poly(I:C) stimulation was used as a positive control. The cytokines in the culture supernatants were quantified by Luminex®. (A) Cytokine concentration in uninfected A2EN cells. Only detectable cytokines are shown. (B) Variation in cytokine expression after CT infection, with or without SP treatment, relative to non-infected (NI) condition. Increase in cytokine

Given the impact of the pool of SP on CT infection, we also tested the impact of individual SP, isolated from patients with different clinical and inflammatory parameters. The individual SP were characterized in terms of cytokine/chemokine concentrations (Figure 4A). The individual SP had various inflammatory profiles, we thus classified them into two groups, one “high inflammation”, with more pro-inflammatory cytokines expressed and another “low inflammation” (Figure 4B). The effect of the individual SP was examined on A2EN cells infected with CT. Several dilutions of SP in PBS were tested. No effect was detected on cell viability. We then determined the percentage of CT infection in the various conditions by quantifying the infected cells by immunofluorescence. We showed that for all the individual SP tested, a 50-fold dilution of SP inhibited the percentage of CT infected cells at 24h post-infection (Figure 4C). This effect was also dose dependent: the percentage of infection increased for a 500-fold dilution of SP. concentration is represented in red, decrease in blue. Asterisks indicate a significant difference between NI and CT or poly(I:C) by one sample t-test (δp≤0.05, δδp≤0.01, δδδp≤0.001) or between CT and SP or CT+SP by two-way ANOVA test (*p ≤0.05, **p ≤0.01, ***p ≤0.001).

**Figure 4:**
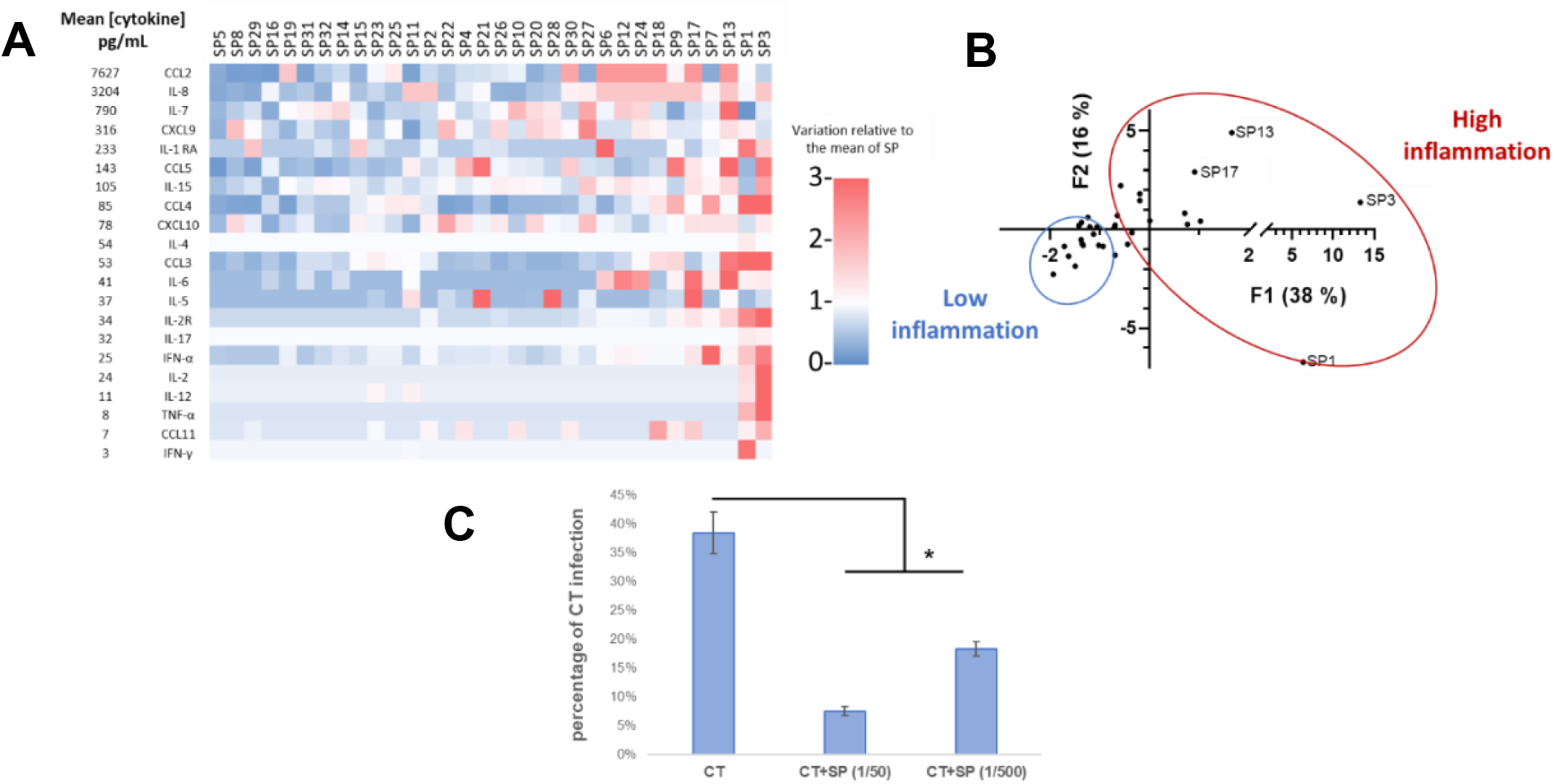
Cytokine concentrations of individual SPs and impact on CT infection of A2EN cells. Cells were infected or not with CT at a MOI of 12, in the presence or not of different dilutions of SP, for 24h (n=2). (A) Cytokine characterization of the individual SPs. The cytokine concentrations were quantified by Luminex® in each of the SP (n=32). (B) A principal component analysis was performed on the cytokine profiles of individual SP, and thus they were classified into two groups, high inflammatory and low inflammatory. (C) The percentage of CT infection was determined by immunofluorescence at 24hpi by quantification of the inclusion in A2EN infected cells. The asterisk indicates a significant difference by T-test (*p ≤0.05).

### Seminal plasma induced transcriptional changes in A2EN cells

To address the effect on gene expression of the inhibition of CT infection of A2EN cells by the pool of SP, a transcriptomic analysis of A2EN cells in various experimental conditions was performed (Figure 5). In CT infected A2EN cells, 162 CT genes are detected at 48h in the transcriptomic profile of A2EN cells. Moreover, 52 eukaryotic genes are differentially regulated following the infection: for example, ATG5, ATG16L1, UBE2I, NEDD4L or many ZNF-genes involved in various metabolic processes and in the regulation of gene expression are upregulated. During exposition to the pool of SP, genes that are statistically differentially regulated are different from those in the non-infected (NI) vs CT infected condition. Some of those genes are metalloproteases (MPST, ADAMTS1). When A2EN cells are infected with CT in presence of the pool of SP, all CT genes are downregulated and only 23 genes are detected at 48h post-infection compared to the NI condition. Interestingly, comparing the host related genes, the cells present a different expression pattern compared to the A2EN cells exposed only to the pool of SP or only to CT. Some genes in the CT vs CT+SP condition are involved in the antigen processing and presentation (NFYC, CTSB). Others are involved in innate immune responses (MIF).

**Figure 5:**
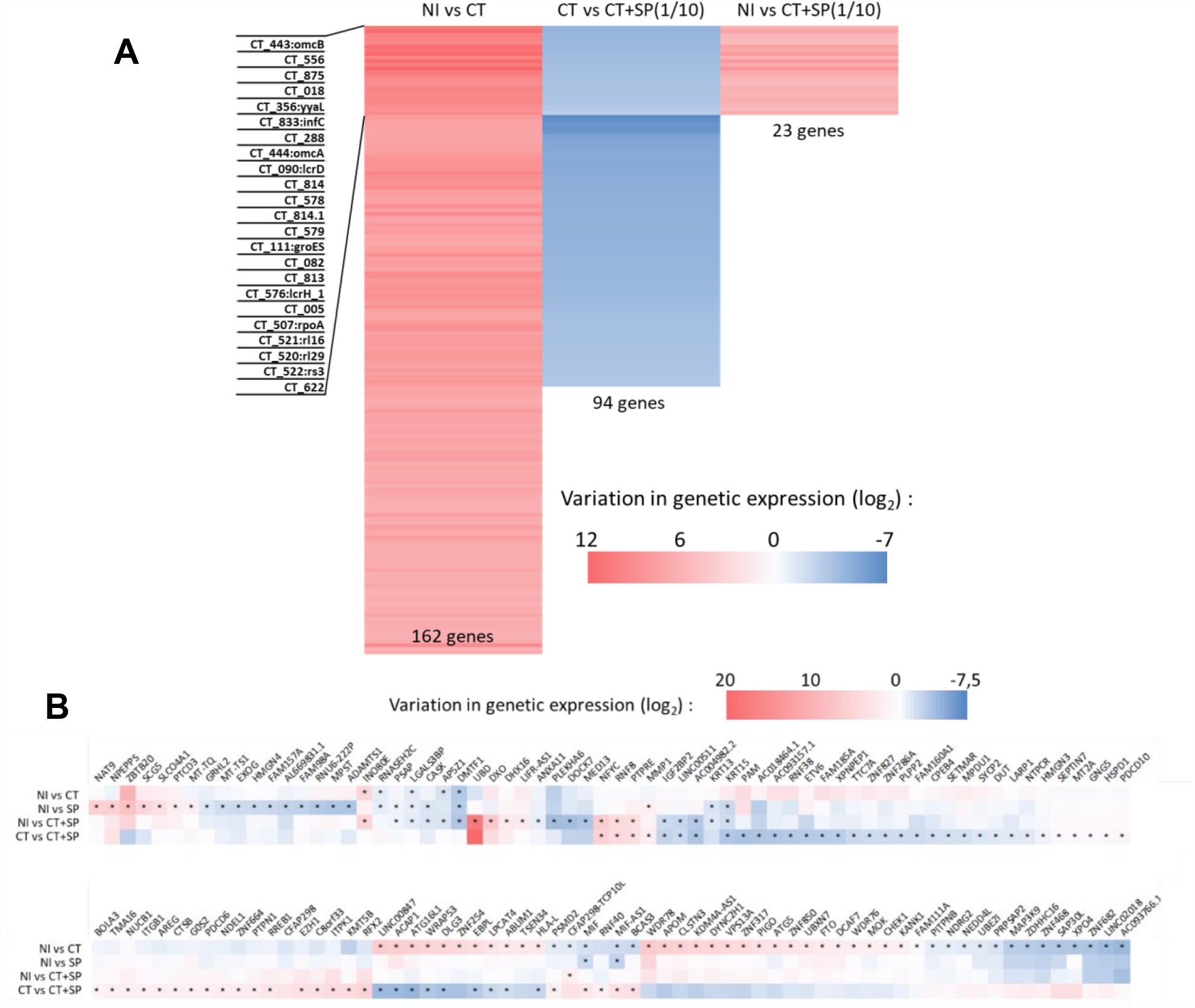
Transcriptomic analysis of A2EN cells infected with CT in presence or not of SP. Cells were infected or not with CT at a MOI of 25 for 48h, in presence or not of a 10-fold dilution of SP and were then lysed for transcriptomic analysis (n=3). *C. trachomatis* (A) or host-responding (B) gene expression in A2EN cells exposed to CT, SP or CT+SP. The log_2_ fold change in gene expression was calculated using Enrichr, and is represented compared either to the uninfected/unstimulated (NI) condition or to CT infected cells (*: adjusted p-value < 0.05).

In summary, CT and SP both induce transcriptional changes in A2EN cells. In A2EN cells infected with CT in presence of SP, the transcription profile observed is even different from the one of the non-infected and/or non-SP exposed cells. CT genes are also affected by the presence of SP.

### Impact of A2EN supernatants on neutrophil phenotype and functions

Given the modifications induced by SP ± CT on epithelial cells, we wondered if the variations in the cytokine concentrations could have an impact on the underlying cells in the mucosa. We focused our study on neutrophils since they are the first cells recruited to the site of inflammation. When recruited, neutrophils mostly come from the blood^31^, so we used neutrophils isolated from human blood to assess the impact of A2EN supernatants on their phenotype and functions.

#### Phenotype

First, neutrophil phenotype was analyzed at 0h and 6h post sorting (Supplementary Figure 4). In DMEM medium, the neutrophil phenotype was comparable between 0h and 6h post sorting, except for a major decrease in CD62L, and a small decrease in CD101 expression. Then, we assessed the impact of A2EN supernatants on neutrophil phenotype after 6h of culture. PD-L1 expression on neutrophils was not affected by A2EN supernatants. All supernatants had a quite similar effect on other cellular markers, with an increase in CD11b, CD32a, CD10, CD101 and CD62L expression on neutrophils (Figure 6). However, the significance threshold was not reached for all conditions: when neutrophils were treated with non-infected and non-exposed to SP (NI) A2EN cell supernatants, there was no significant increase in CD10. Moreover, CT infected A2EN cell supernatants did not significantly delay the degradation of CD62L. The A2EN supernatants resulted in at least a 2.5-fold increase in the mean fluorescence intensity (MFI) of CD11b, CD32a and CD62L, whereas the increase in the MFI of CD10 and CD101 was less important and did not exceed 100%. Finally, the decrease in CD62L expression observed in neutrophils after 6h of culture was less drastic when neutrophils were treated with A2EN supernatants, and more particularly with supernatants from A2EN cells treated with a 10-fold dilution of SP. In those conditions, the MFI was at least 7 times more important compared to the DMEM condition. The difference in CD62L MFI was significantly different between the NI and SP (1/10) conditions, but it was only a tendency (p=0,2) between the NI and CT+SP (1/10) conditions. In conclusion, in all the tested conditions, A2EN supernatants increased the expression of several surface markers in blood neutrophils.

**Figure 6:**
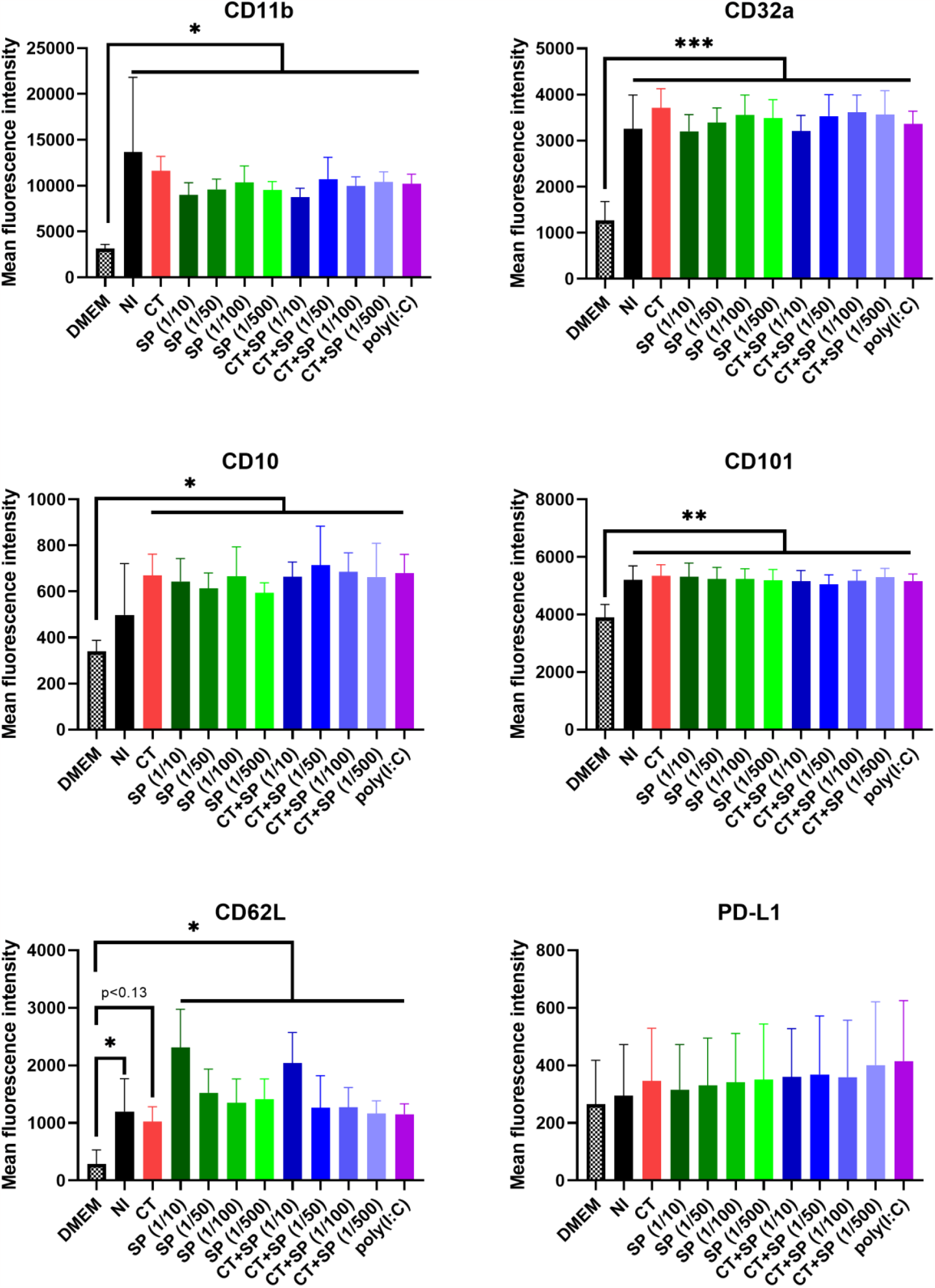
Impact of the A2EN cell supernatants on blood neutrophil phenotype. A2EN cells were infected or not with CT at a MOI of 12, in the presence or not of different dilutions of the pool of SP (n=4). Poly(I:C) was used as a positive control. After 24h, the supernatants were collected and used on neutrophils isolated from the blood. Neutrophil phenotype was then analyzed after 6h of incubation, using antibodies listed in Table 1. The mean fluorescence intensity of CD11b, CD32a, CD10, CD101, CD62L and PD-L1 was analysed on neutrophils (CD45^+^ CD66^+^ Lin (CD3, 8, 14, 20, 123, 125). Asterisks indicate a significant difference by two-way ANOVA test (*p ≤0.05, **p ≤0.01, ***p ≤0.001).

#### Survival

Neutrophil survival was evaluated at 2h, 6h, 24h and 48h post sorting (Figure 7A). In DMEM medium, the survival of the neutrophils was drastically reduced as soon as 6h post sorting and continued to diminish at 24h, with 60% and 20% of cell survival respectively. In contrast, A2EN supernatants enhanced significantly neutrophil survival after 6h of culture (p<0.01), with at least 13% and 40% of cell survival at 6h and 24h respectively. More precisely, at 24h of culture, supernatants of A2EN cells infected with CT promote neutrophil survival (67.3% of neutrophil survival) compared to the conditions without CT (at best 51.7% of neutrophil survival) (Figure 7B). SP also had an effect because supernatants from CT infected A2EN cells, treated with various dilutions of SP, affected the neutrophil survival in a dose dependant manner. Indeed, while SP dilution increased, CT infected A2EN supernatants increased the survival of the neutrophils (from 40.6% of neutrophil survival in the 1/10 dilution to 57.5% of neutrophil survival in the 1/500 dilution). Altogether, when treated with SP, the effect of CT infected A2EN supernatants was dependent on the SP dilution: at high SP dilutions, the effect of CT predominated with an enhanced survival of the neutrophils comparing to low dilutions (Figure 7B).

**Figure 7:**
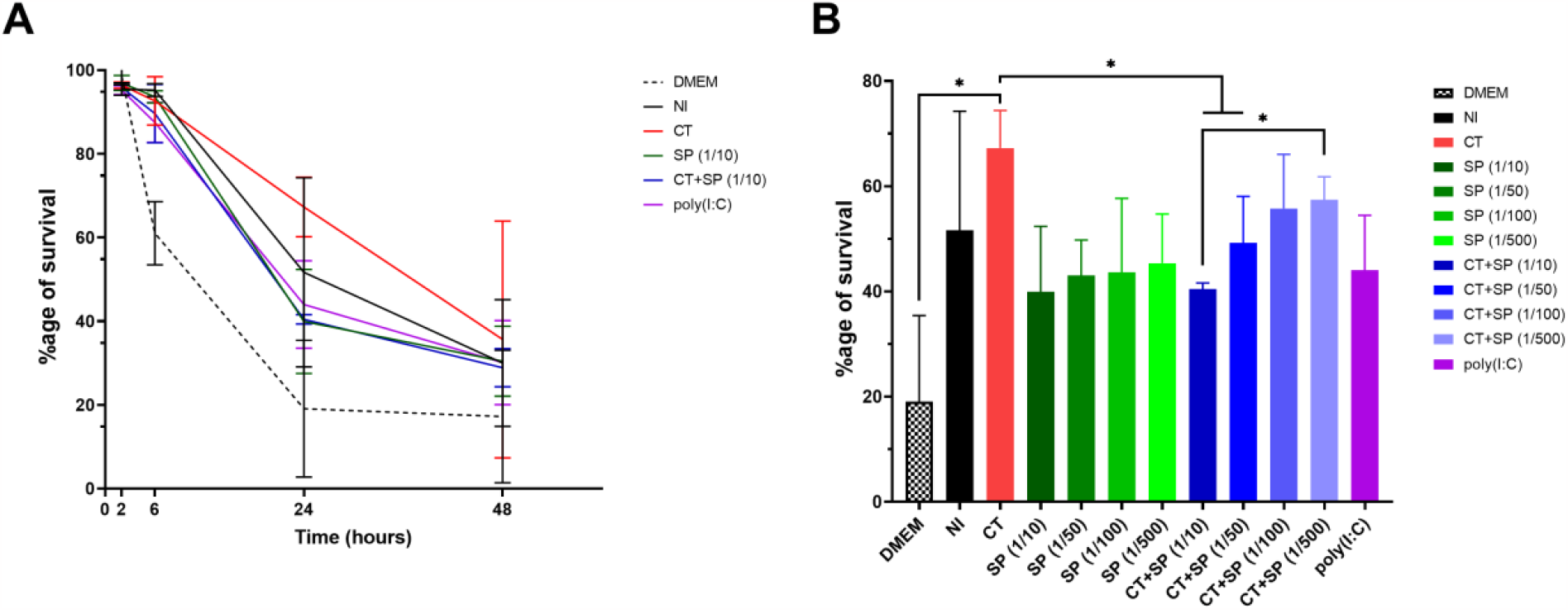
Impact of the A2EN cell supernatants on blood neutrophil survival. A2EN cells were infected or not with CT at a MOI of 12, in the presence or not of different dilutions of the pool of SP (n=4). Poly(I:C) was used as a positive control. After 24h, the supernatants were collected and used on neutrophils isolated from the blood. Neutrophil survival was then studied after 2h to 48h of incubation, using AnV and 7-AAD antibodies. Live cells (AnV^-^ 7-AAD^-^) were quantified among neutrophils using FlowJo software. (A) Kinetics of the neutrophil survival. Results are expressed as a percentage of survival. (B) Variation of neutrophil survival after 24h of incubation with the different A2EN supernatants. Asterisks indicate a significant difference by two-way ANOVA test (*p ≤0.05).

#### ROS production

Neutrophils were incubated 2h with DMEM medium or A2EN supernatants, and ROS production was evaluated by stimulating neutrophils with PMA and monitoring the emission of light by oxidized luminol. A representative graph of the 90min follow-up is illustrated Figure 8A and Figure 8C. When a PMA stimulation was performed at t=0h (Figure 8A and 8B), neutrophils cultured in DMEM medium were less susceptible to PMA induced ROS production than neutrophils in A2EN treated with SP or poly(I:C) supernatants (Figure 8B). Moreover, in DMEM medium, the peak of ROS production occurred a few minutes later than for neutrophils in A2EN supernatants (Supplementary Figure 5). Without any PMA stimulation, ROS production by neutrophils was very weak and did not differ between the experimental conditions, but neutrophils were still able to produce ROS upon later PMA stimulation (Figure 8C). After 50 min in PBS, PMA stimulation induced a ROS production by neutrophils in all experimental conditions. Neutrophils cultured in DMEM medium tended to induce less ROS production than neutrophils in A2EN treated with SP or poly(I:C) supernatants, even if the difference was not significant (Figure 8D). CT infection did not significantly impact the effect of A2EN supernatants on ROS production. In conclusion, poly(I:C) and SP treated A2EN cell supernatants are potent activators of PMA induced ROS production in neutrophils.

**Figure 8:**
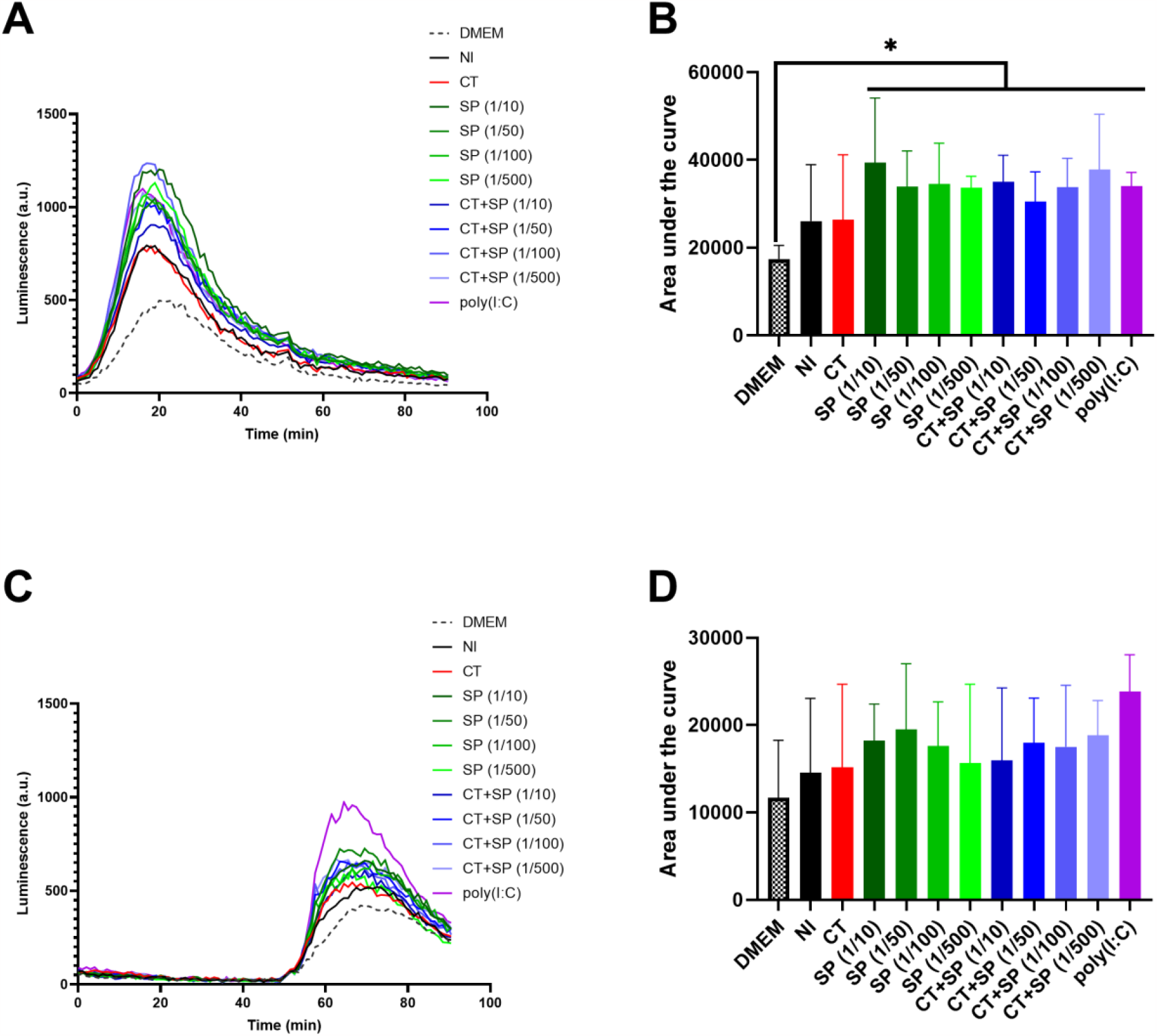
Impact of the A2EN cell supernatants on blood neutrophil ROS production. A2EN cells were infected or not with CT at a MOI of 12, in the presence or not of different dilutions of the pool of SP (n=4). Poly(I:C) was used as a positive control. After 24h, the supernatants were collected and used on neutrophils isolated from the blood. After 2h of incubation, ROS production was evaluated for 90min by monitoring luminescence emitted by oxidized luminol. PMA was added to stimulated ROS production, either directly at t_0_ (A, B) or after 50min of luminescence monitoring (C, D). A representative read-out of the experiment is provided in both cases (A, C). The area under the curve is also represented in both cases (B, D). Asterisks indicate a significant difference by two-way ANOVA test (*p ≤0.05).

#### Phagocytosis

Neutrophil phagocytic capacity was evaluated at 2h post sorting, at 37°C and 4°C. The MFI of neutrophils was evaluated to assess their phagocytic capacity (Figure 9A). The MFI for all experimental conditions at 4°C was lower than at 37°C (MFI_max 4°C_= 57 whereas MFI_min 37°C_= 138). The minimal MFI at 37°C was obtained for neutrophils cultured in DMEM medium, and 58% of neutrophils were engaged in the phagocytosis process (data not shown). When they were cultured in A2EN supernatants, the phagocytosis activity was increased and at least 83% of neutrophils were engaged in the phagocytosis process (data not shown). The MFI of the neutrophils in uninfected A2EN supernatants was significantly lower than for the other A2EN supernatants, meaning that their phagocytic activity was significantly increased (Figure 9B). Indeed, supernatants of stimulated (by poly(I:C), CT and/or SP) A2EN cells caused an increase of at least 70% of neutrophil phagocytic capacity compared to unstimulated A2EN supernatants. In conclusion, A2EN cell supernatants, and more particularly stimulated A2EN cell supernatants are potent activators of phagocytic capacity in neutrophils.

**Figure 9:**
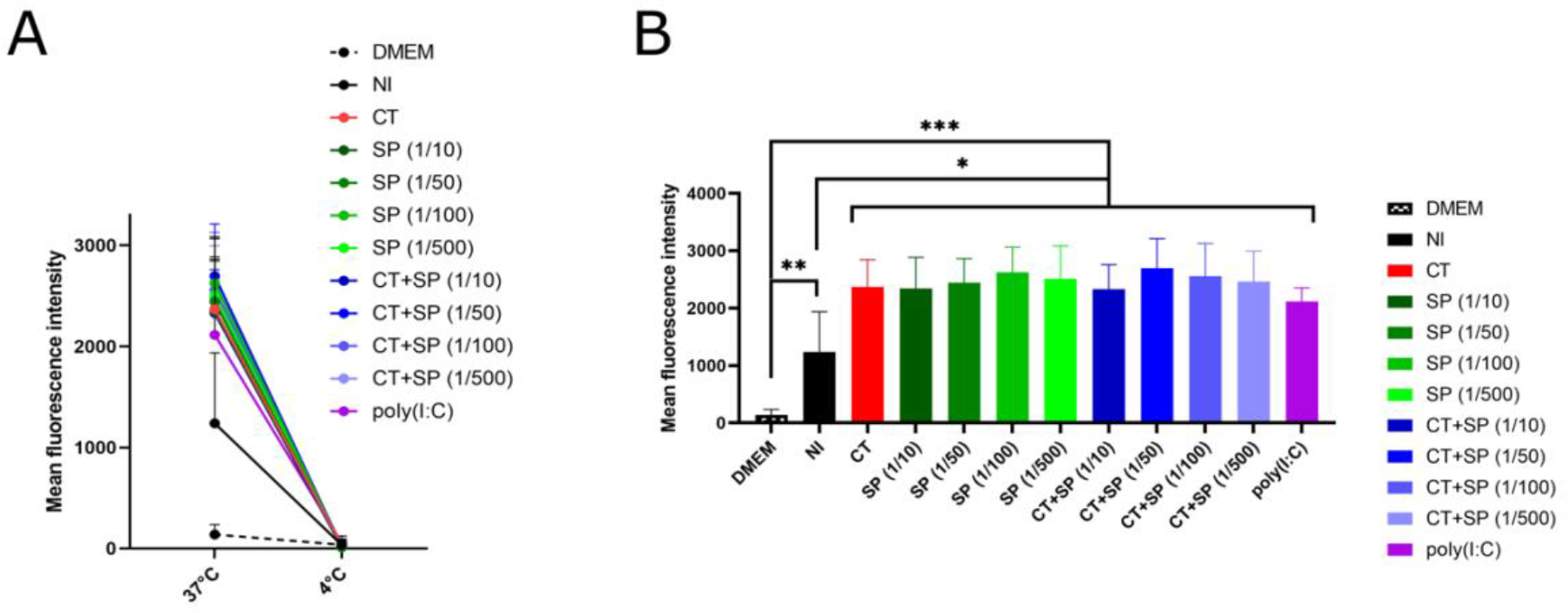
Impact of the A2EN cell supernatants on blood neutrophil phagocytosis capacity. A2EN cells were infected or not with CT at a MOI of 12, in the presence or not of different dilutions of SP (n=4). Poly(I:C) was used as a positive control. After 24h, the supernatants were collected and used on neutrophils isolated from the blood. After 2h of incubation, fluorescent *E. coli* particles were added to the neutrophils for 5min, cells were then fixed and the mean intracellular fluorescence was assessed by flow cytometry. (A) Comparison of the mean fluorescence intensity reflecting the phagocytic capacity in the different experimental conditions at 37°C and 4°C. (B) Impact of A2EN supernatants on neutrophils at 37°C. Asterisks indicate a significant difference by two-way ANOVA test compared to the baseline (*p ≤0.05, **p ≤0.01, ***p ≤0.001).

## Discussion

The environment of the FRT, and more particularly the inflammation, is particularly important for the transmission of STI. However, the underlying mechanisms are often poorly or not documented, particularly in the context of co-infections^32^. During this study, we developed an *in vitro* model aiming at better understanding the role of the SP and the mucosal environment (in terms of inflammation) during CT infection. This model allows to study sequentially the different steps occurring at the level of the FRT *in vivo* during CT infection: cervicovaginal epithelial cells are infected by CT in presence of SP, then the epithelial cells secrete inflammatory mediators which act on immune cells in the *lamina propria* ^6^.

*In vitro* and *in vivo* studies have shown that SP is a potent immunomodulator and affects STI acquisition, either positively or negatively^33^. During male to female transmission of CT, SP acts as a vehicle for the bacteria. In this study, we reported for the first time that SP inhibits CT svD infection of human endocervical epithelial cells. We did not find any correlation between the percentage of CT infection in the different conditions and the cytokine profiles of the individual SP used. The analysis of the transcriptional profile of A2EN cells infected with CT in presence or not of SP revealed that CT genes were more frequently detected in absence of SP. The bacterial genes expressed in the CT and CT+SP conditions were comparable with those found in a previous study of CT infection in HeLa cells^34^. We observed a reduction of many immediate early genes (CT_288, CT_111:groES, CT_529, CT_446:euo, CT_774:cysQ, …) expressed in the CT+SP condition compared to the CT condition. This could suggest a delay in the CT growth in presence of SP, however, we did not find any difference in the size of the inclusion at 24h post-infection, suggesting an inhibition of early stages of the CT infection, which may include entry stages. The infection inhibition by SP may thus be at several steps: factors in the SP could affect the entry of the bacteria leading to a reduction of the percentage of infection at 24h and at later stages of the infection, the presence of SP could affect the bacterial growth within the cells at 48h post-infection. This multi-step model is compatible with the CT life cycle since CT complete its cycle in 48h to 72h^35^. Several host genes were also differentially regulated by SP and/or CT. However, we were not able to find general pathways implicated during SP and/or CT exposition. The factor(s) in the SP impacting CT infection do not appear to be the cytokines/chemokines that we have quantified, since all the individual SP tested, with various inflammatory profiles, had the same impact on the percentage of CT infection, suggesting that another factor may be involved. This factor might be another cytokine, that we did not investigate in this study, or defensins for example, that have been described to be present in SP and have been shown to have antimicrobial properties^36^.

A2EN cells infected with CT produce pro-inflammatory cytokines at 24 hpi, but the increase in cytokine concentration was relatively modest compared to poly(I:C) stimulation, as already reported in the literature^5^. For example, IL-8 concentration was not increased by CT or by poly(I:C), or in any other of the experimental conditions. Buckner et al., showed that it was possible to stimulate the production of IL-8 by A2EN cells using poly(I:C), but these results were obtained in different experimental conditions, and with polarized cells. Moreover, the fold increase in the cytokine concentration was correlated with the percentage of CT infection. Also, the basal level of IL-8 was already very high at baseline in our experiments, as it has been reported at the level of TRF^37^. This high level of IL-8 has been associated with the recruitment and activation of immune cells^6^.

On the other hand, SP had an anti-inflammatory effect on epithelial cells. The pool of SP we used had a moderate inflammatory profile, and TGB-β was the most concentrated cytokine. This cytokine could be the main modulator of A2EN cytokine production, since it is a known modulator of cytokine production in other cervical cell lines^38^. It has been involved in numerous immunomodulatory processes, notably in accommodating the foreign alloantigens to promote immune tolerance to favor a successful pregnancy^39^. Nevertheless, the anti-inflammatory properties of the SP observed in this study are in contrast to what has been described after a SP inoculation in the FRT mucosa *in vivo*^39^. This difference can be explained because the *in vitro* model does not take into account the complexity of the FRT, which contains a diversity of cells (including resident immune cells) that could be responsible for the pro-inflammatory reaction to SP. The cytokines found in the supernatants of A2EN cells infected with CT with SP seem to confirm the anti-inflammatory effect of SP on epithelial cells: the presence of SP leads to modifications in the expression of cytokines by CT-induced A2EN cells. These modifications may be due either to the fact that SP interfere with CT infection cycle, and/or to an additive effect: the expression profile obtained corresponds to the sum of the two profiles obtained with the SP and CT.

We then tested the impact of A2EN supernatants on neutrophils isolated from the blood, since those cells are rapidly recruited after CT infection^6^. However, neutrophils were directly isolated from the peripheral blood, and thus their phenotype may differ from extravasated neutrophils located *in vivo* at the site of CT infection. We showed that A2EN supernatants affect the phenotype of neutrophils, increasing the level of CD11b and CD32a on the one hand, suggesting a priming of neutrophils and increasing the levels of CD10 and CD101 on the other hand, suggesting a maturation of neutrophils^40^. For most surface markers, there was no difference according to the A2EN supernatants used, so this argues in favor that those phenotypes could be associated to highly concentrated cytokines in all A2EN supernatants such as IL-8, also found in high concentration at the level of the FRT, and that has an impact on neutrophil *in vivo*. The increase in CD62L, when neutrophils were exposed with SP treated A2EN supernatants, may indicate a delay in CD62L degradation by A2EN supernatants. CD62L is a L-selectin involved and cleaved during diapedesis. This delay in CD62L degradation by A2EN supernatants can be correlated with the survival of the neutrophils. We showed that A2EN supernatants increase neutrophil survival, suggesting that the soluble factors in A2EN supernatants have a globally positive effect on neutrophil survival. This could be due to IL-6 or more probably GM-CSF, because those two cytokines are strongly decreased in SP treated A2EN supernatants. Indeed, GM-CSF has been shown to impact neutrophil surface marker expression^41^: for example, a GM-CSF stimulation has been shown to reduce CD62L expression at the level of neutrophils or eosinophils^42,43^. Regarding the functions of neutrophils, the effect of A2EN supernatants was heterogeneous according to the function studied. For the survival assay, the effect of A2EN supernatants was in line with their cytokine profile: each condition had a different effect, with CT infected cells conferring the highest neutrophil survival. For CT infected cells, the effect of SP also depended on the dilution: at high dilutions, the effect of CT dominated with an enhanced neutrophil survival compared to lower dilutions. We can thus argue that the cytokines that are differentially expressed in the supernatants of A2EN cells infected with CT (namely CCL5, CCL3, CXCL10, IL-6 or GM-CSF) may impact neutrophil survival. Concerning phagocytosis and ROS production, all A2EN supernatants increased these functions in neutrophils, suggesting that highly concentrated cytokines in all A2EN supernatants (such as IL-8), could impact neutrophil phagocytosis and ROS production. Indeed, IL-8 has been shown to be able to prime neutrophils^44^, in presence of a second stimulus such as LPS, neutrophil activation can trigger phagocytosis and ROS production^45^. We also demonstrated that the supernatants of non-infected A2EN cells had a lower impact on neutrophil phagocytic activity compared to the other A2EN cell supernatants. This would suggest that the stimulation of A2EN cells create an environment that promotes the phagocytosis of neutrophils.

We showed that SP could impact the inflammation at the level of the FRT during CT infection. Those modifications could also impact the risk for STI acquisition and lead to co-infections, so we also tested the impact of A2EN cell supernatants on the susceptibility to HIV-1 infection. However, we were not able to highlight significant differences in U87-CD4-CCR5 susceptibility to a virus pseudotyped with HIV-1 R5 envelope or with VSV-G protein (data not shown). This model allowed to assess the effect of A2EN supernatants in a simplified system with a one cycle viral replication and to evaluate if the supernatant’s impact was occurring at the step of the viral entry. However, it does not recapitulate the full diversity of the immune cells present at the level of the FRT, nor the complexity of a HIV-1 replicative clinical isolate. It will be interesting to test A2EN cell supernatants on primary cells isolated from the FRT, and infected with a HIV-1 R5-tropic virus. Overall, our results highlight the need to take into account SP when studying the mechanisms of STI acquisition. SP has a significant impact on the susceptibility to CT infection and on the innate immune responses to the infection. The interactions between the SP and other factors from the environment at the level of the FRT should be taken into consideration to develop new preventive strategies against STI such as CT.

## Acknowledgments

The authors would like to thank L2I team (Julie Morin, Laetitia Bossevot) in the IDMIT department for their help to perform the multiplex assay for cytokine detection, as well as members of the Life&Soft company (Ségolène Diry, Léo d’Agata, Cassandra Gaspar and Eric Ginoux) for the transcriptomic experiments. The authors would also like to thank Dr Frank Follmann (SSI, Danemark) for kindly providing the CT strain used for the *in vitro* infection of A2EN cells, Drs Luc de Chaisemartin (UMR 996, Hôpital Bichat, France) and Julien Lemaître (IDMIT department, CEA, France) for providing advice concerning the development of the neutrophil functional assays and Dr Agathe Subtil (Institut Pasteur, Cellular Biology of Microbial infection unit) for helpful discussion on CT infection and inhibition.

The authors thank all the patients who provided samples, AP-HP (Assistance Publique Hôpitaux de Paris) and clinical personnel for collecting the samples; the Clinical Core of the CRT (Center of Translational Science) of the Institut Pasteur for their help with biomedical regulatory aspects of the project, in particular Anaïs Perilhou.

