## Supplemental figures for "Seminal plasma inhibits Chlamydia trachomatis infection *in vitro*, and may have consequences on mucosal immunity"

Supplementary Figures

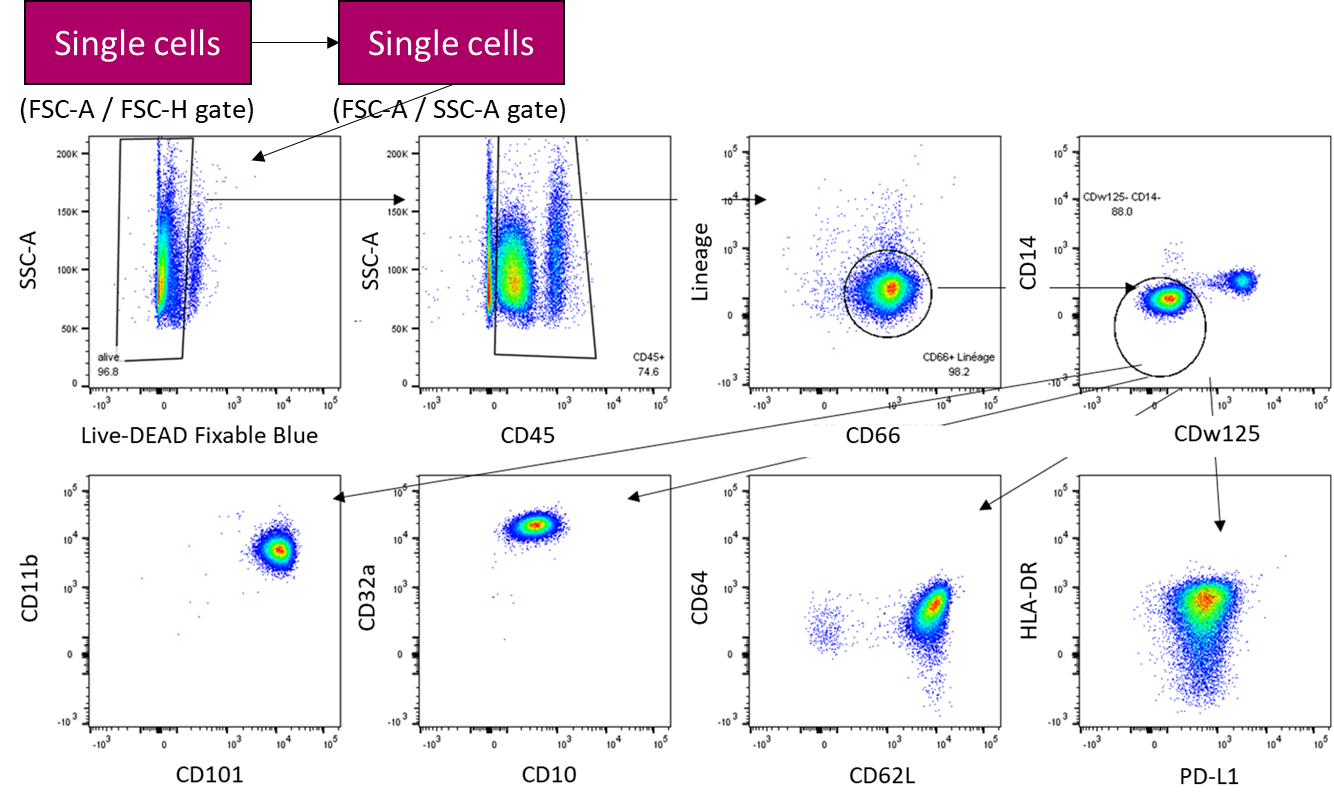

**Supplementary Figure 1: FACS gating strategy for blood neutrophil phenotyping**. Neutrophils were defined as CD45^+^, CD66^+^, Lin (CD3,CD8,CD20,CD123)^-^, CD14^-^, CDw125^-^ cells. The mean fluorescence intensity (MFI) of CD11b, CD101, CD10, CD32a, CD62L and PD-L1 was then determined. The MFI of CD64 and HLA-DR were very low and did not vary throughout the study.

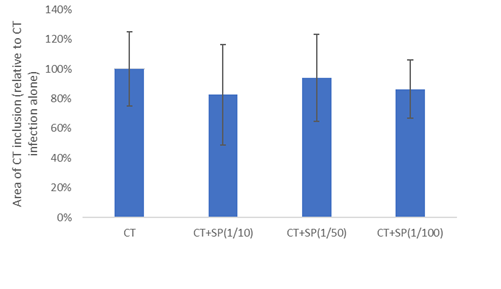

**Supplementary Figure 2: Impact of SP on CT inclusion area**. A2EN cells were infected with CT at a MOI of 12 for 24h in presence or not of different dilutions of the pool of SP (n=3). The size of the CT inclusions was determined by quantification of the area of CT inclusion in A2EN infected cells by immunofluorescence staining. The results are expressed relatively to the CT infection without SP.

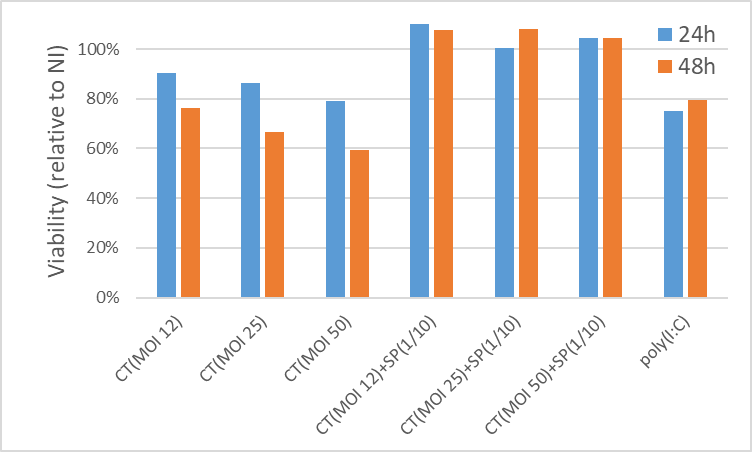

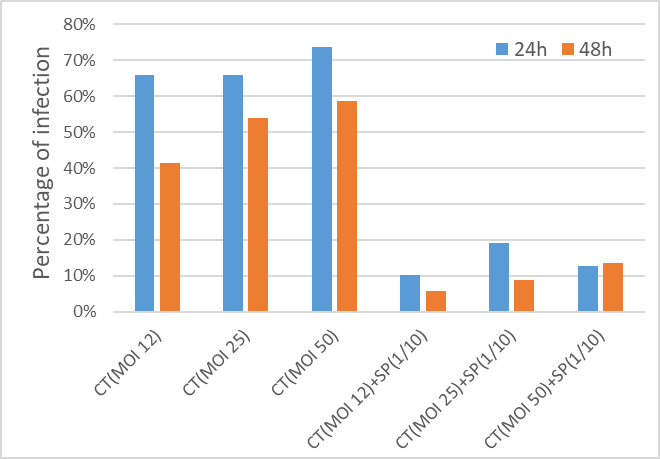
**Supplementary Figure 3: Impact of SP on CT infection at different MOI**. A2EN cells were infected with CT at a MOI of 12, 25 or 50 for 24h or 48h (n=1). (A) A2EN viability was evaluated using CellTiter. (B) The percentage of infection was determined by quantification of the CT inclusion in A2EN infected cells by immunofluorescence staining.

B

A

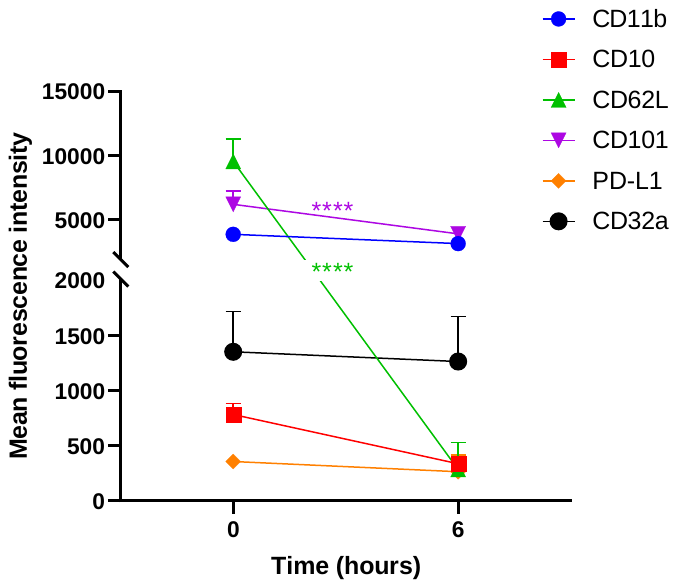

**Supplementary Figure 4: Variation of the MFI of neutrophil markers.** Neutrophils were isolated from human blood and cultured in 50% DMEM + 50% RPMI for 6h (n=4). Their phenotype was studied after 0h and 6h of incubation, using the antibodies listed in Table 1. The mean fluorescence intensity of CD11b, CD32a, CD10, CD101, CD62L and PD-L1 was analyzed in neutrophils. Asterisks indicate a significant difference by one-way ANOVA test (****p ≤0.0001).

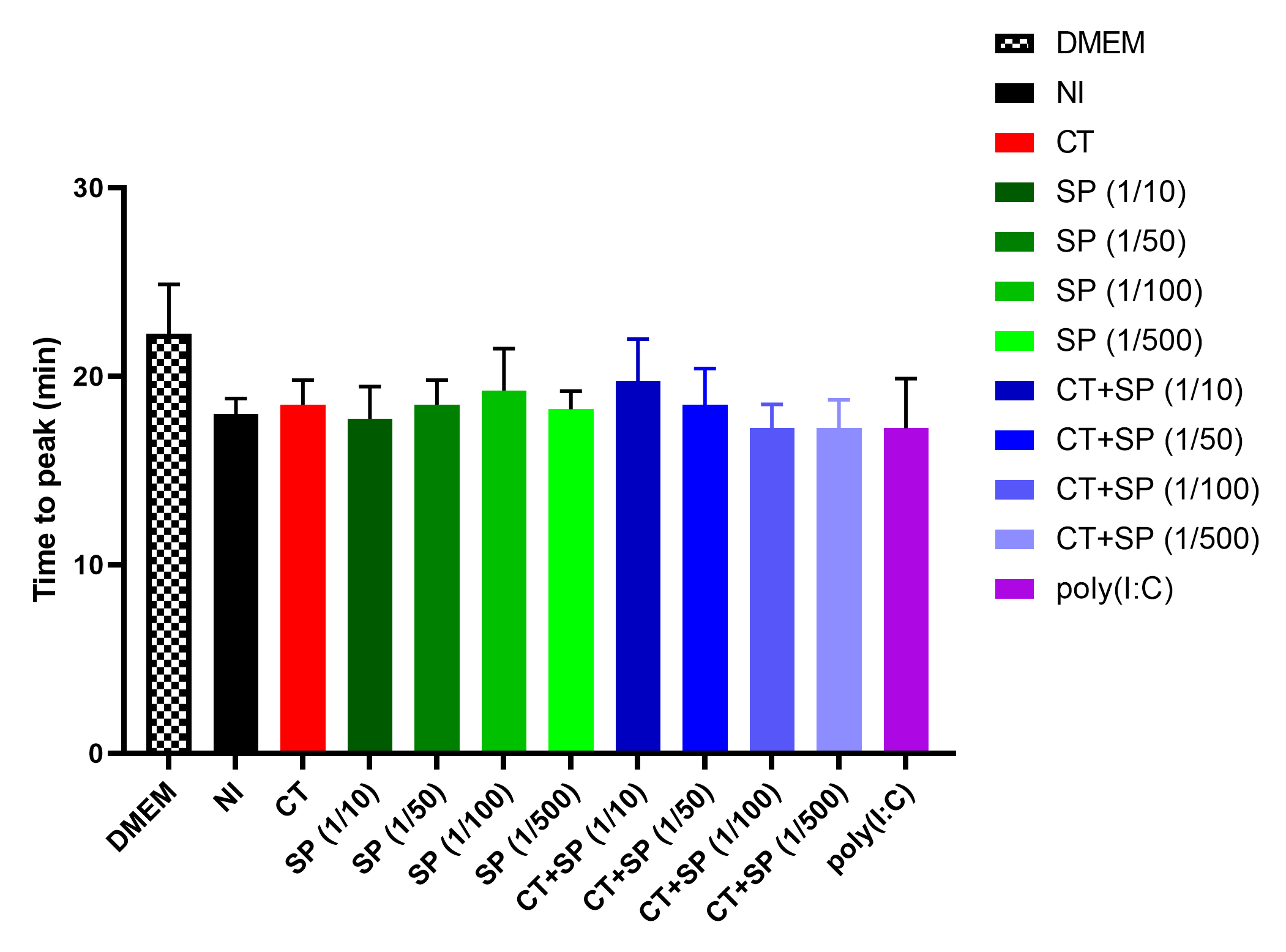

**Supplementary Figure 5:** **Impact of** **A2EN cell supernatants on blood neutrophil ROS production.** A2EN cells were infected or not with CT at a MOI of 12 in presence or not of different dilutions of the pool of SP (n=4). After 24h, the supernatants were collected and used on neutrophils isolated from the blood. After 2h of incubation, PMA was added and ROS production was evaluated for 90min by monitoring luminescence emitted by oxidized luminol. The time to reach the peak of ROS production is shown here.
